# Bladder pressure encoding by near-independent fibre subpopulations — implications for decoding

**DOI:** 10.1101/826297

**Authors:** Carl H. Lubba, Zhonghua Ouyang, Nick S. Jones, Tim M. Bruns, Simon R. Schultz

**Author notes:** These authors contributed equally to this study (Tim M. Bruns, Simon R. Schultz).

## Abstract

**Objective:** We aim at characterising the encoding of bladder pressure (intravesical pressure) by a population of sensory fibres. This research is motivated by the possibility to restore bladder function in elderly patients or after spinal cord injury using implanted devices, so called bioelectronic medicines. For these devices, nerve-based estimation of intravesical pressure can enable a personalized and on-demand stimulation paradigm, which will be more effective and efficient. In this context, a better understanding of the encoding strategies employed by the body might in the future be exploited by informed decoding algorithms that enable a precise and robust bladder-pressure estimation.

**Approach:** To this end, we apply information theory to microelectrode-array recordings from the cat sacral dorsal root ganglion while filling the bladder, conduct surrogate data studies to augment the data we have, and finally decode pressure in a simple informed approach.

**Main results:** We find an encoding scheme by different main bladder neuron types that we divide into three response types (slow tonic, phasic, and derivative fibres). We show that an encoding by different bladder neuron types, each represented by multiple cells, offers reliability through within-type redundancy and high information rates through near-independence of different types. Our subsequent decoding study shows a potentially more robust decoding from mean responses of homogeneous cell pools.

**Significance:** We have here, for the first time, analysed the encoding of intravesical pressure by a population of sensory neurons in a principled way using information theory. We show that even a simple adapted decoder can exploit the redundancy in the population to be more robust against cell loss. This work thus paves the way towards principled encoding studies in the periphery and towards a new generation of informed peripheral nerve decoders for bioelectronic medicines.

## 1. Introduction

A new paradigm for the treatment of diverse medical conditions has recently emerged: the development of ‘bioelectronic medicines’ [1] which modulate the peripheral nerve signaling by means of implanted devices. This alternative to molecular medicine has promise as a localized and permanent remedy for conditions as varied as hypertension and tachycardia [2, 3], sleep apnea [4], rheumatoid arthritis [5] and many others. Most current neuromodulation devices still operate in a simple open-loop fashion, acting in a preset way, independent of changes in the physiological processes they try to influence. In the future, bioelectronic medicines are expected to become more advanced and include real-time feedback about current organ states. By only blocking or stimulating when necessary, closed-loop devices can be much more efficient and effective, and even capable of dynamically managing conditions, e.g., detecting parasympathetic bronchoconstriction in asthma and suppressing it [6].

To enable the use of feedback control, physiological quantities of interest may be measured by chemical, mechanical, or other sensors that are implanted in addition to the nerve interface [7, 8, 9, 10, 11, 12]. While this approach seems straightforward from an engineering point of view, surgery becomes more difficult and the probability of complications (e.g., device movement, tissue damage, loss of signal [7, 13]) post surgery rises. An alternative approach is to harness, where possible, the body’s own sensors for monitoring and control of organs. Thousands of afferent fibres continuously transmit signals about organ physiological state. These existing biological bladder neurons are sensitive and may offer a stable source of organ state information as an elegant alternative to implanted artificial sensors. In order to take full advantage of these signals, however, we need a better understanding of how physiological quantities are encoded by populations of peripheral afferent fibres. We can then implement informed decoders tailored to the encoding strategies present in the periphery, and use them as a robust and precise feedback in next generation bioelectronic medicines.

The bladder provides an ideal testbed for the development of closed-loop bioelectronic medicines. It has a main parameter of interest – its fullness, characterized by both volume and resulting intravesical pressure – which can easily be manipulated and recorded. The bladder wall is further covered by numerous stretch sensors that monitor this central quantity. It is thus a good candidate for investigating the encoding of an organ parameter by a multitude of cells. Developing closed-loop bioelectronic medicines for the bladder is furthermore clinically important, as bladder dysfunction is a common condition both in the elderly population [14], and after spinal cord injury [15, 16]. The resulting incontinence has devastating effects on a patient’s quality of life [17, 18]. In addition, other malfunctions such as detrusor-sphincter dyssynergia and hyper-reflexia can occur in specific patient groups and cause renal damage, repeated urinary tract inflammations and kidney infections [19, 20].

The lower urinary tract (LUT), consisting of the bladder, urethra and sphincter, is innervated by the pelvic, the pudendal, and the hypogastric nerves [21]. The pelvic nerve projects to the internal pelvic organs including bladder, urethra, bowel, and vagina [22, 23]. The pudendal nerve goes to the pelvic floor including urethra, sphincter, anal sphincter, perineal region, genitalia [24, 25, 26]. The hypogastric nerve forms a plexus with the pelvic nerve, innervating similar regions, including the bladder neck/proximal urethra. We are therefore mainly interested in pelvic nerve fibres that originate in the sacral-level dorsal root ganglion (DRG) to innervate the bladder wall. In the cat, most cell bodies giving rise to the afferent fibres projecting through the pelvic nerve to the bladder can be found in sacral-level S1 and S2 DRG [27, 28]. Of the approximately 40000 cell bodies in the cat sacral DRG S1 and S2 [29], about 1000 innervate the bladder [30, 28, 31, 32]. This population is composed of both small myelinated Aδ and unmyelinated C-fibres, of which the former are generally accepted to transport the mechanoreceptor impulses and trigger the normal micturition reflex [21, 33, 34, 35]. C-fibres are associated with nociception but have been reported to sense bladder pressure (intravesical pressure) in addition to Aδ fibres [36]. The bladder neuron responses were characterized as tonic (Aδ) and phasic (C-fibres) [37], sometimes described as pressure (Aδ) and volume (C) receptors [38] and are usually found to have a diversity of activation thresholds within each diameter range [39, 40, 41, 36]. Some exhibit hysteresis [42]. While the large body of physiological studies draws a detailed descriptive picture of bladder afferents, elucidating the physiological significance of the different cell types for pressure encoding has not been attempted. The question of why the diverse bladder neuron responses exist is one we seek to answer in this work.

Just as physiologists have led a rich variety of studies on the afferent innervation of the bladder, engineers explored various ways to decode intravesical pressure, volume and contractions from peripheral nerve activity in the past. Intravesical pressure is the most informative quantity, as all other events like for example the onset of (reflexive) detrusor contractions can be detected from the pressure time course. Decoders using pelvic [43, 13, 44], pudendal [45, 46] or spinal nerves [47, 48] have been proposed. Targeting these nerves, however, requires a difficult surgery and recordings using the common cuff interface often lack good signal-to-noise ratio (SNR) without severely damaging the nerve [47]. As an alternative, one can interface with sacral-level dorsal root ganglion (DRG) where cell bodies of both pelvic and pudendal nerve fibres reside. Recording cell bodies with penetrating microelectrode arrays (MEA) is more invasive than the cuff electrode, but leads to a good signal-to-noise ratio at high spatial resolution. The development of axon-sized electrodes (microns or tens of microns dimensions) will minimize immune responses as a result of electrode insertion, making this approach feasible despite its high invasiveness. Moreover, the activation of efferent pathways can be accomplished at the same site through reflex circuits [49, 50]. Decoding from microelectrode arrays implanted in the DRG has been demonstrated [39, 51, 52], with a stable interface over weeks [50]. While many of the above decoding approaches, be it from a peripheral nerve or from the DRG, estimate intravesical pressure quite accurately, none of them directly draw on insights from physiological studies of the encoding. Most proposed solutions rely on single cell responses (if spatial resolution and SNR allow) and are assumed to be stable over time. If a change in the recording setup occurs, however, e.g., due to electrode migration, cell death, etc, the decoder has no means of detecting this change, and can quickly lose its prediction quality without retraining.

In the present work, we investigate both the encoding of pressure across the many afferent fibres innervating the bladder wall, and draw conclusions towards building better decoders that exploit the observed encoding strategies. We conducted this research based on microelectrode array recordings from the sacral dorsal root ganglion levels S1 and S2 in cats (see Fig. 1A for the experimental apparatus) during a slow filling at a near physiological rate. We distinguished three distinct stereotypical response types that recur across experiments: slow tonic, phasic, and derivative. For each type, we used information theory to quantify the information it individually carries about intravesical pressure, and further estimated the benefits of combining different bladder neuron types – on both real and simulated data. Taking advantage of the insights gained from this information theoretic encoding analysis, we propose an informed decoding strategy from stereotypical groups of fibres that proves to be robust and accurate.

**Figure 1:**
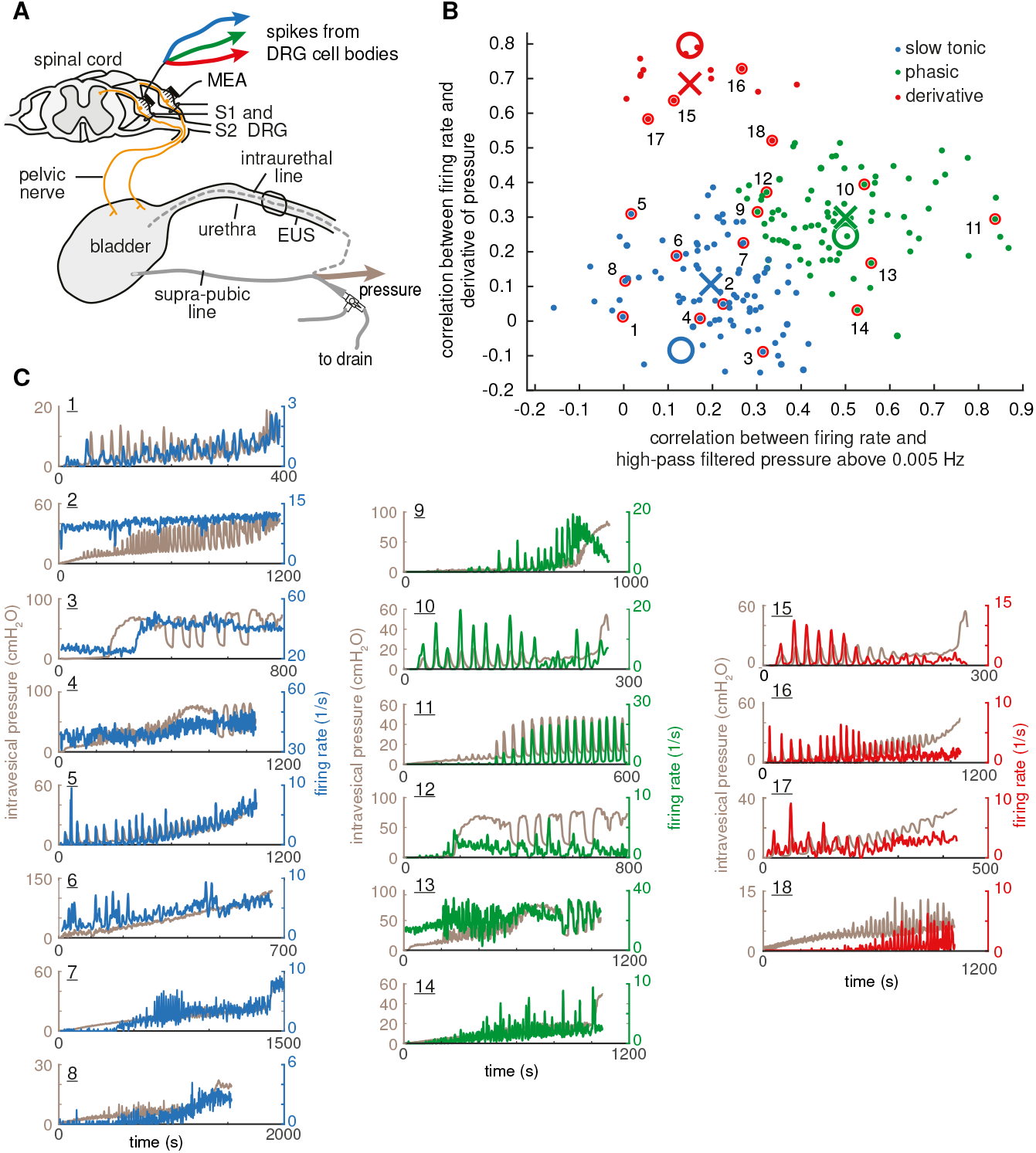
Intravesical pressure is encoded by distinct groups of stereotypical cells. **A** Microelectrode array (MEA) recording of cells from the first and second sacral dorsal root ganglion (DRG S1/S2) along with the intravesical pressure. The bladder was filled through a catheter in the bladder dome (solid line) or urethra (dotted line). Graphic adapted from [42]. **B** When plotting all fibres of all trials in the 2D-plane of the correlations of firing rates with high-pass filtered pressure (x-axis) and derivative of pressure (y-axis), we can associate regions of this correlation-feature plane with the different bladder neuron types shown in (B). Crosses indicate the cluster centers obtained through k-means clustering and large circles show the manually selected initial centers. **C** Example firing rates of bladder neurons along with the corresponding intravesical pressure are shown for different trials. By the indices the plots can be related to the scatter in panel B. The cells stem from the following experiments and trials (E: Experiment, T: trial, C: cell index). 1: E1 T57 C1, 2: E3 T100 C15, 3: E4 T28 C22, 4: E4 T29 C23, 5: E5 T57 C5, 6: E6 T24 C1, 7: E7 T19 C2, 8: E8 T68 C7, 9: E2 T9 C1, 10: E2 T11 C7, 11: E3 T74 C10, 12: E4 T28 C12, 13: E4 T29 C3, 14: E7 T18 C1, 15: E2 T11 C2, 16: E5 T57 C1, 17: E5 T58 C7, 18: E8 T67 C1.

## 2. Methods

### 2.1. Experiments

We analyse here data previously collected for a study of single-unit hysteresis [42] and a comparison of intravesical pressure decoding algorithms [51] (experiments 1-5) and a study on real-time decoding of intravesical pressure [52] (experiments 6-8). Full details of experimental procedures can be found in those respective publications. In short, 8 adult male cats of approximately 1 year of age were used. All procedures were approved by the University of Michigan Institutional Animal Care and Use Committee, in accordance with the National Institute of Health guidelines for the care and use of laboratory animals. For experiments 1 and 5 a 5×10 microelectrode array (Blackrock Microsystems, Salt Lake City, Utah, USA) was inserted in the left S1 DRG and a 4×10 microelectrode array into the left S2 DRG. For experiments 2, 3 and 4, 5×10 arrays were inserted bilaterally in S1 and 4× 10 arrays were inserted bilaterally in S2. Experiments 6 to 8 had 4× 8 electrode arrays in left and right S1. Microelectrode shank lengths were either 0.5 or 1.0 mm with 0.4 mm inter-shank spacing. Intravesical pressure was recorded simultaneously with a catheter either inserted through the urethra or inserted into the bladder dome, at a sampling rate of 1 kHz and low-pass filtered for further analysis at 4 Hz.

The experimental apparatus is shown in Fig. 1A. We emptied the bladder using the bladder catheter before filling it with saline at a near-physiological rate of 2 ml/min [53]. This filling cystometry rate was just above the maximal physiological rate typically reported for felines (up to 15 x 1.1 ml/kg/hr, which is 1.1–1.7 ml/min for the animals in this study) [54]. Inflow was stopped when we observed dripping from the external meatus or, if present, around the urethral catheter. The saline had room-temperature (22°C) in experiments 1-4 and 6–8 and body-temperature (41°C) in experiment 5. Two filling cystometry trials per experiment with only non-voiding bladder contractions form the basis of the following analysis (without the final voiding phase). Trials took 17 min on average (minimum 5 min, maximum 30 min). Neural signals were recorded at 30 kHz with a Neural Interface Processor (Ripple LLC, Salt Lake City, Utah).

After data collection, voltage signals on each microelectrode channel had an amplitude threshold between 20 and 35 μV applied (3–5.5 times the rootmean-square of the signal) to identify spike snippets of neuron action potential firings. Spike snippets were sorted conservatively by experienced researchers in Offline Sorter v3.3.5 (Plexon, Dallas, TX), using principal component analysis, followed by manual review to identify unique spike clusters. In MATLAB (Mathworks, Natick, MA), instantaneous firing rates for each cell were then calculated by smoothing with a non-causal triangular kernel [55] of width 3 s.

### 2.2. Fibre selection and characterization

We first inspected fibre responses manually. In this process, we observed different but recurring response characteristics across experiments. Some cells would respond with a gradual change to rising pressure, and some would mainly respond to quick changes and decay into inactivity for phases of constant pressure, even if high. Among the fibres responding to quick components, some seemed to respond to positive changes in pressure rather than reacting to the raw pressure and then adapting. From this manual inspection of the responses that were all different from each other, we defined three main distinct response types, (1) ‘slow tonic’: a mainly monotonic rise in firing rate with mean pressure across long time scales without coverage of the quick non-voiding contractions, (2) adapting ‘phasic’ fibres which respond to quick changes in intravesical pressure during contractions but, because they adapt over time, do not report the mean pressure with the same fidelity as ‘slow tonic’ ones, and (3) ‘derivative’ fibres which only respond to phases of rising pressure and are, similar to phasic fibres, weakly indicative of the mean pressure. The firing rates of derivative fibres have a similar appearance to phasic fibres, but in fact their firing rates were shifted in time with respect to each other, the derivative fibres reacting earlier than the phasic fibres.

In a next step, we sought to associate the cells of our experiments automatically to these three types across all experiments. As pressure time courses varied from trial to trial, we could not directly compare the firing rate time courses of the bladder neurons across different trials. We therefore devised features that described the reaction of each neuron to the intravesical pressure. The three features we selected were the Pearson correlation coefficients between firing rates and (1) low-pass filtered pressure below 0.01 Hz, (2) high-pass filtered pressure above 0.005 Hz, and (3) derivative of pressure ‡ for every fibre of every trial. The high- and low-pass filter cutoff frequencies were chosen to separate the pressure signal into a slow mean component without contractions and the contractions only. As a first step of our automated selection and typing procedure, we only considered neurons as bladder units that had a raw correlation *ρ* above 0.4 to at least one of these filtered intravesical pressure variants. This is an arbitrary cut-off that we selected based on experience from previous decoding studies where including weakly correlated fibres did not improve decoding performance [51]. As our experiments contained many candidate cells (~1000) to choose from, we could afford an increased selectivity to be sure to only consider unambiguously relevant neurons.

We obtained the two-dimensional plane of the correlation measures between firing rate and high-pass filtered pressure (x-axis) and derivative of pressure (y-axis), shown in Fig. 1B. The 2D-plane of the two high-frequency correlation features gave us all the relevant information, because we had already removed cells without correlation to any of the three pressure signal variants. Thus, cells with low correlations to both derivative and high-pass filtered pressure were implicitly identified as only highly correlated to the slow component. In the scatter plot of Fig. 1B, the fibres occupy different regions.

To associate bladder neurons globally across all trials with the three response types we identified manually, we chose to conduct a *k*-means clustering with *k* set to 3 in in this ‘correlation-feature’ plane. The converged and initial centers are shown in Fig. 1B. This approach will be referred to as ‘feature clustering’ and forms the basis of associating each cell to one of the three bladder neuron types slow tonic, phasic, and derivative.

We emphasize that the clustering in feature-plane does not claim to uncover number and characteristics of fibre types in a purely data-driven, unsupervised clustering approach. It is part of a semi-automatic cell typing in which each bladder neuron is associated to one of the three types we identified manually in a reproducible, objective manner. The semi-automatic nature of our method is especially obvious for the lower cluster that we separated into slow tonic and phasic fibres where no clear separation is visible. We could have united the two parts into a single cluster. However, while some cells exhibited mixed responses, many of them clearly corresponded to either a phasic or a slow tonic behavior. See Fig. 1C for examples. Therefore, reducing to two clusters would have combined neurons of clearly different response characteristics to a single type. As the constraints of our experimental data (number of neurons, recording length, and repetitions) did not allow us to divide the cells fully automatically, we chose to keep some degree of subjectivity, trusted the initial manual cell typing, and split the large lower cluster into the two types slow tonic and phasic. Any overlap in response characteristics was interpreted as noise. We note that this separation of cells into three types is a simplification that may (1) misclassify cells, especially the ones of mixed type and (2) shadow more subtle differences within pseudo-homogeneous types. However, we believe classification is essential, to isolate independent information. As a side note, no finer frequency analysis was possible, as the pressure signals did not contain spectral power above approximately 0.2 Hz.

Clustering by correlation-features relied on the availability of the pressure signal and its differently filtered variants. As an unsupervised alternative that could, for instance, be carried out on-line in an implanted device, we also clustered fibres hierarchically in each trial based on the pairwise Pearson correlation coefficients between their firing rates (number of clusters fixed to the number of fibre types in each trial). This step allowed us to identify clusters of similarly evolving firing rates – and therefore fibres of similar bladder neuron characteristics – without relying on the intravesical pressure. We will denote this approach ‘activity clustering’. Clustering by activity was only possible within each trial, not across trials, as pressure dynamics differed between trials.

### 2.3. Surrogate data

The dataset had three inconvenient properties which affected the analyses: (1) the low-frequency power of the pressure signal was correlated with the high-frequency power as non-voiding contractions mostly occur at high pressures, (2) not all cell types we distinguished were present in all experiments, and (3) the recording length complicated the estimation of information theoretic quantities (detailed in the next section). To partially overcome these limitations, we created surrogate cells (single cells or populations of similar cells) that approached the behavior of the main bladder neuron types, and drove them by idealized stimuli (‘pressure signals’). A surrogate cell consisted of an ‘intended firing’ rate with was derived from the pressure signal (e.g., a low-pass filtered version of the pressure signal) that defined, together with an intended mean firing rate, the rate parameter of an inhomogeneous Poisson process to generate a spike train. See Table 1 for a list of the implemented fibre response types. We define a theoretical ‘tonic’ fibre whose intended firing rate perfectly matches the intravesical pressure and a theoretical ‘linear’ fibre that rises linearly with time (coefficients *a* and *b* fit to data). The remaining three types of simulated cells match the response characteristics we found experimentally. The ‘slow’ fibres follow the low-pass filtered pressure at 0.0005 Hz, ‘derivative’ cells were driven by the pressure derivative, and ‘phasic’ responses were defined using a decay parameter τ in seconds that regulates how quickly the fibre adapts. From the spike times output by the inhomogeneous Poisson process, we computed the continuous firing rate just as we did in the real data by a non-causal triangular kernel of width 3 s.

**Table 1:** Surrogate fibre responses in relation to a stimulus *s*(*t’*). The operation \**h*_LP_ indicates convolution with a low-pass filter.

| Fibre type | response formula |
| --- | --- |
| Tonic | $r(t') = s(t')$ |
| Linear | $r(t') = a + b \cdot t'$ |
| Slow | $r(t') = s(t') * h_{LP}$ |
| Derivative | $r(t') = \frac{ds(t')}{dt}$ |
| Phasic | $\frac{dr(t')}{dt} = \max(\frac{ds(t')}{dt}, 0) - \frac{1}{\tau} r(t')$ |

### 2.4. Information theoretic analysis

Information theory [56, 57], originally developed for the study of communication channels in engineered systems, has proven to be a useful tool in neuroscience for quantifying the information carried by a single cell or a population of neurons about a variable of interest [58, 59, 60, 61, 62]. We here consider a common information theoretic quantity, the Shannon mutual information (MI), estimating (1) how much information each fibre carries about the pressure stimulus, and (2) how much benefit there is in combining the information from two different fibres or fibre types. In the continuous case, mutual information *I*(*X,Y*) is computed between two variables *X* and *Y* of probability distributions *PX* (*x*) and *PY* (*y*) and joint distribution *P* (*x,y*). In our case, *X* could for instance be the firing rate of a selected cell and *Y* could be the intravesical pressure signal. *I*(*X,Y*) then quantifies the amount of entropy of variable *X* that is lost when knowing what values *Y* assumes in all joint measurements of *X* and *Y*:

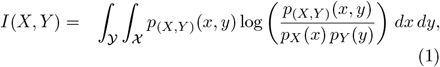

In addition to the two-variable case, we can also quantify the joint mutual information that two variables *X* and *Y* carry together about a third variable of interest *Z*:

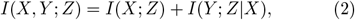

where for *I*(*Y; Z | X*) we have to adapt Eq. 1 by replacing all distributions of *X* and *Y* by conditionals to *Z* and integrate over the distribution of *Z*. We can further combine the individual mutual information measures of both *X* and *Y* and their joint mutual information about *Z* to a quantity called ‘fractional redundancy’, 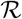, which can assume values between −1 and 1 and indicates how much less information the ensemble of *X* and *Y* contains about *Z* than the sum of the parts,

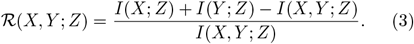

Note that negative values of redundancy imply synergistic interaction between variables. We compute the described information theoretic quantities from firing rate and pressure at a sampling rate of 1/s using the Kraskov mutual information estimator for continuous signals [63], implemented in the JIDT toolbox [64] which we run from MATLAB (Mathworks, Natick, MA). The conditional mutual information needed for the joint MI (Eq. 2) was computed in the full joint space [65, 66] as implemented by the JIDT toolbox.

As our trials were of limited length (1036 ± 399 samples), mutual information estimates were upwardly biased due to finite sampling effects, which are incompletely removed by the Kraskov estimator. This is illustrated in Fig. 2 for a single and a pair of simulated tonic fibre(s). At 1000 samples, mutual information of a single fibre was overestimated by approximately 7%, joint mutual information by approximately 14%, and redundancy was underestimated by 12%. From 10 000 samples, joint and single mutual information as well as the redundancy stabilized.

**Figure 2:**
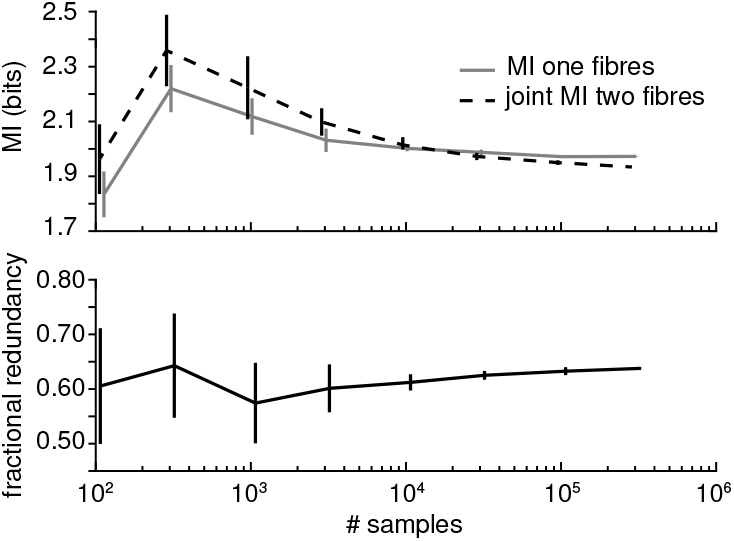
Finite sampling bias results in mild overestimation of mutual information and joint mutual information, but a slight underestimation of redundancy. Mutual information, joint mutual information and redundancy were computed from the firing rate of a simulated tonic fibre (pair) and an idealized pressure signal. Firing rate was set to 20/s for single fibres, and 10/s for each fibre in a pair; 5 repetitions for each signal length. See Sec. 2.3 and Fig. 6 for details of the simulated data.

### 2.5. Decoding

So far we described the quantification of information that individual fibres and fibre combinations carry about pressure - on both real and surrogate data. Using the following approach, we made use of our refined understanding of the physiological encoding and designed an adapted decoder. When estimating intravesical pressure from nervous activity, we face two main areas of choice to be made. (1) The pre-processing of the neural signal: whether we sort cells or take some measure of activity per electrode, what cells we choose if sorted, how we compute the spike rate, and (2) the type of decoding algorithm we use: Optimal Linear Estimator (OLE), Kalman filter, neural networks, etc. We focus here on the pre-processing based on the sorted cell responses and fix the decoding algorithm to an OLE [67] and a Kalman filter [68] for comparison. For the optimal linear filter, the decoded pressure 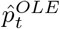 is then obtained from the firing rates vector ***f_t_*** and the regression coefficients **β** at a given time t:

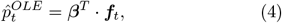

where **β** minimizes the mean-squared error *E* (*N* number of time-points):

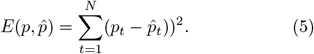

From our encoding results, we compare three signal variants to decode from in both estimation error and robustness against cell loss - a common problem due to electrode migration or cell death:

1. all single sorted cells
2. cells of each fibre type pooled (as in Fig. 1B)
3. cells of each activity cluster pooled

For pooled signal variants, we normalized the firing rate of each cell by its mean to make sure their contributions were added with equal weight. We tested the robustness of these different pre-processing variants to cell loss by training on all fibres, and removing a randomly chosen 20% of the cells before testing. As a measure of decoding quality, we computed the normalised root mean squared error (NRMSE) between decoded pressure 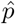 and true pressure *p* as in Eq. 6 with *p_min/max_* minimum and maximum pressure and *N* number of samples. All errors were averaged across 5 cross-validation folds within triels.

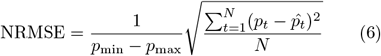

To compare decoding performances statistically, we conducted paired *t*-test across trials. The statistical tests were repeated with only the first trial of each experiment to exclude that the independence of observations was violated.

## 3. Results

We found 185 bladder-units within 1044 overall fibres across 22 trials in 8 animals by thresholding the Pearson correlation coefficient between firing rates and the pressure signals (see Sec. 2.2 for details). We treat these 185 fibres as separate from each other even though some will have been recorded repeatedly across multiple trials of the same experiments.

### 3.1. Groups of stereotypical bladder neuron types exist

As shown in Fig. 1B and described in Sec. 2.2, in order to associate each cell with one of the three types we defined, we first clustered cells globally by the correlation of their firing rates to the high-pass filtered pressure and the pressure derivative (‘correlation-features’). In this way, we distinguished 89 cells as slow tonic, 81 as phasic and 15 as derivative. The derivative cells were clearly separated from the other types in correlation-feature plane (Fig. 1B). It is noteworthy that 12 of 15 derivative cells stem from the two trials of one experiment. Tonic and phasic showed a more gradual transition. Some slow tonic fibres also responded to quick contractions to some extent, and some phasic fibres did not completely decay to inactivity for stimulus plateau phases. In addition to these overlapping receptor properties, the stimulus signal did not separate phases of high low-frequency power and high high-frequency power well, as non-voiding contractions mostly occurred in the high-pressure regime. In many trials, this caused the firing rates of quick phasic fibres to be correlated with the slow component of the pressure as well. It is thus to be expected that some cells were misclassified as a certain type. We further reiterate that the separation into three distinct types is a deliberate simplification. An overview of the cell types in each trial is given in Table 2. Each trial was usually dominated by one or two fibre type(s).

**Table 2:**
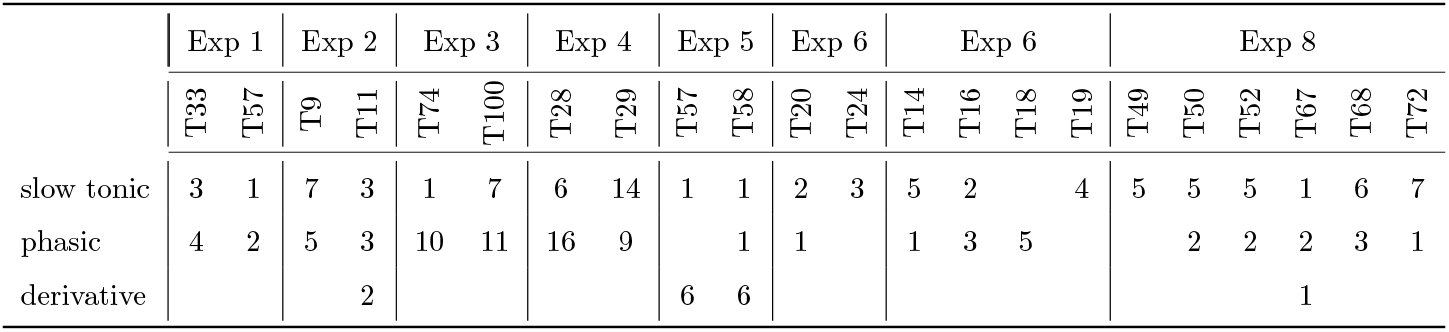
Summary of the identified bladder units across trials. Numbers are fibre counts, TXX indicates experiment-specific trial numbers, among trials for other objectives in each experiment.

The clustering described above required knowledge of the pressure signal in order to compute the correlation-features. As an alternative, we attempted to retrieve the cell types in an unsupervised way by grouping similarly firing cells within each trial to activity clusters. Because similar response characteristics should produce similar outputs given the same stimulus, the activity clusters should correspond to feature clusters (cell types) we observed across all trials. As Table 3 shows, activity clusters often reproduce cell types well. We here assigned a cell type label to each cluster from the dominant type. Fig. 3 shows an example of an activity clustered trial with a clean separation of fibre types. In the normalized single neuron firing rate traces in Fig. 3B the diversity of cell responses within each type, especially within the slow tonic type, is visible. Table A1 gives a more detailed overview of the relation between activity clusters and cell types in all trials.

**Figure 3:**
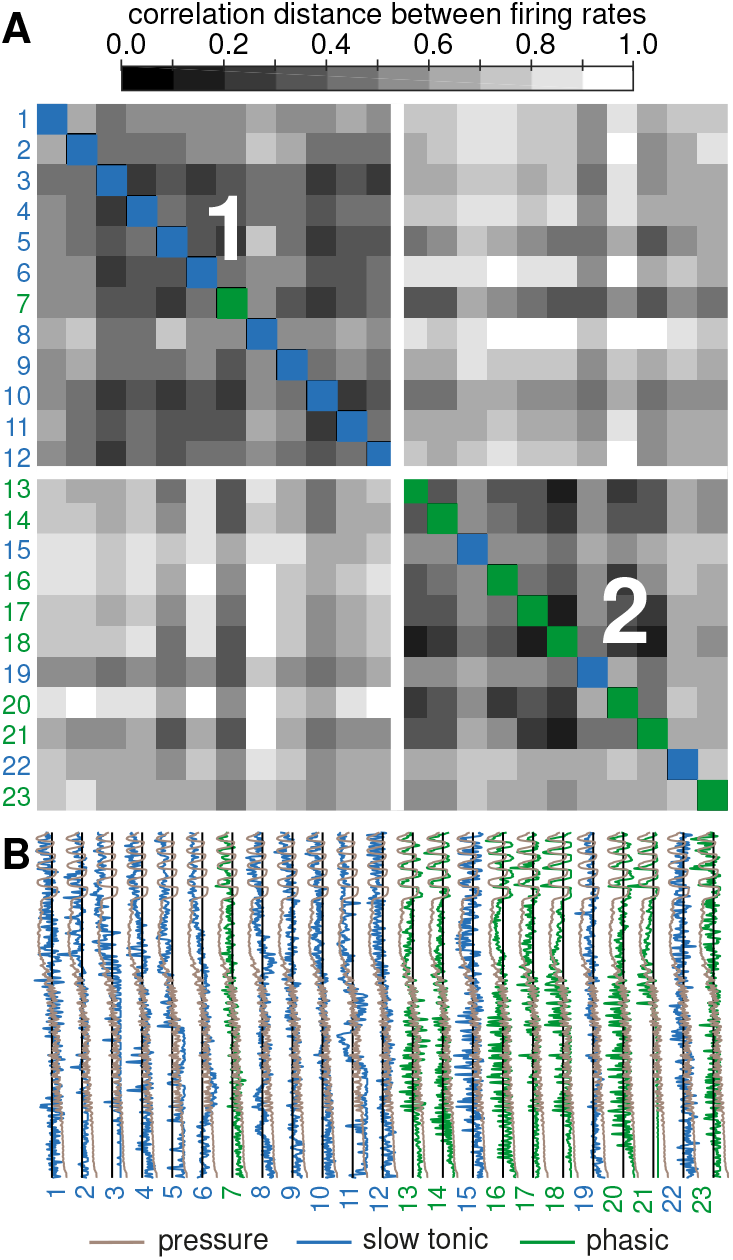
Activity clusters per trial correspond to bladder neuron types; example trial E4T29. **A** The correlation distance matrix shows the two main clusters. Overall, both clusters are quite homogeneous in their cell type content (see colored squares). **B** The time course of all normalized bladder unit firing rates along with the intravesical pressure.

The presence of imperfectly tuned fibres that respond to both static pressure and to quick pressure changes complicated a clean clustering into bladder neuron types, particularly a clean distinction of the types phasic and tonic. Also, slow tonic fibres could have low pairwise similarity complicating the activity clustering. We still often retrieved the same fibre groups by both global clustering across all trials based on correlation-features and by simply grouping similarly firing fibres per trial. It was thus feasible to cluster cells online into different bladder neuron groups by their activity.

The bladder neuron types were not localized at distinct electrodes within the MEA. Most often, one electrode recorded from one neuron only. If multiple cells were recorded by a single electrode, these were rarely of a single type. Fig. 4 shows two example maps of two experiments where many electrodes recorded from multiple cells and Table 4 lists the share of the dominant fibre type per mixed electrode (multiple neurons). E.g., if an electrode records from 3 cells, the dominant share can be either 1 (3/3) or 0.667 (2/3) or 0.333 (1/3). Because fibre types were not uniformly distributed within trials, we also provide the shares after randomly shuffling the electrode types, showing what values would be expected if cell types were distributed randomly across the electrodes. The dominant shares of the real data werechance consistent, meaning that there was no grouping of bladder neuron types within the MEAs. Still, the cell types per electrode were quite homogeneous because often the entire trial was dominated by one fibre type.

**Figure 4:**
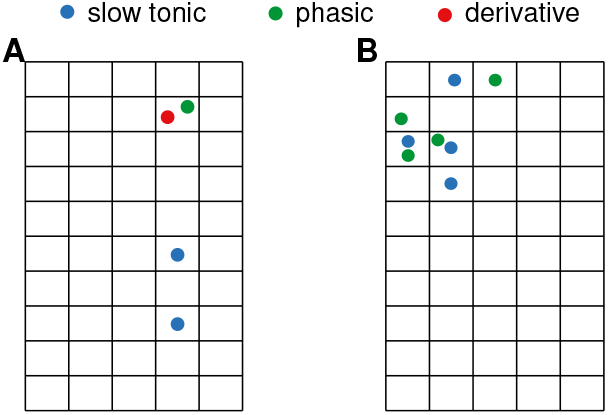
Example distribution of cell types on the MEAs. **A** Experiment 2, trial 11. Right S1 DRG. **B** Experiment 4, trial 29. Right S1 DRG. Microelectrode arrays with most cells shown.

**Table 3:** Clustering by activity within trial often recovers the cell types obtained from clustering in correlation feature plane. Rows show the dominant fibre type in each activity cluster, columns give the fibre type identities from correlation feature plane clustering.

|  |  | Fibre types contained in cluster |  |  |
| --- | --- | --- | --- | --- |
|  |  | slow tonic | phasic | derivative |
| Dominant fibre type in cluster | slow tonic | 74 | 15 | 0 |
|  | phasic | 14 | 65 | 1 |
|  | derivative | 1 | 1 | 14 |

**Table 4:** Cell types only cluster at electrodes to a chance consistent degree. Electrodes recorded from up to 3 bladder neurons. The values are the shares of the dominant fibre type within each electrode across trials (mean ± standard deviation). The shuffled values give an estimation of what share would be expected if the cell types were randomly distributed across the electrodes (cell types shuffled 10 times).

| # neurons | share dominant type | shuffled |
| --- | --- | --- |
| 1 (N=120) | $1.000 \pm 0.000$ | $1.000 \pm 0.000$ |
| 2 (N=25) | $0.820 \pm 0.245$ | $0.794 \pm 0.134$ |
| 3 (N=5) | $0.800 \pm 0.183$ | $0.733 \pm 0.033$ |

### 3.2. Encoding by groups of stereotypical bladder neurons is efficient and robust

We have shown that different but recurring bladder neuron responses exist exist in the studied animals and separated them into three types. In the following section, we aim to identify reasons for both the observed response diversity (different types) and the presence of multiple very similar bladder neurons.

Fibres of the same type were highly redundant, as indicated by the diagonal elements of Fig. 5A. § A straightforward way of making use of this redundancy and quantifying the benefit of duplicating sensors is to pool these fibres into a single compound activity signal. Such pooling of similar sensors enhances the mutual information: the MI of the averaged firing rates of one fibre type on the diagonal of Fig. 5B is substantially higher (at least by a factor of 4, often more) than the average *single* fibre mutual information shown on the diagonal of Fig. 5C (according to the small values the heatmap appears very dark, see Table B2 and Table B3 for all MI values). As single units map intravesical pressure (or an aspect of it such as the slow rise) imperfectly due to both their tuning (e.g., activation threshold) and the spiking nature of their output, pooling many similar cells increases the information content. The signal-to-noise ratio is enhanced through averaging many imperfect sensor outputs [69]. Fig. 5E further illustrates the benefit of averaging over multiple redundant cells: information rises with fibre count in almost every case.

**Figure 5:**
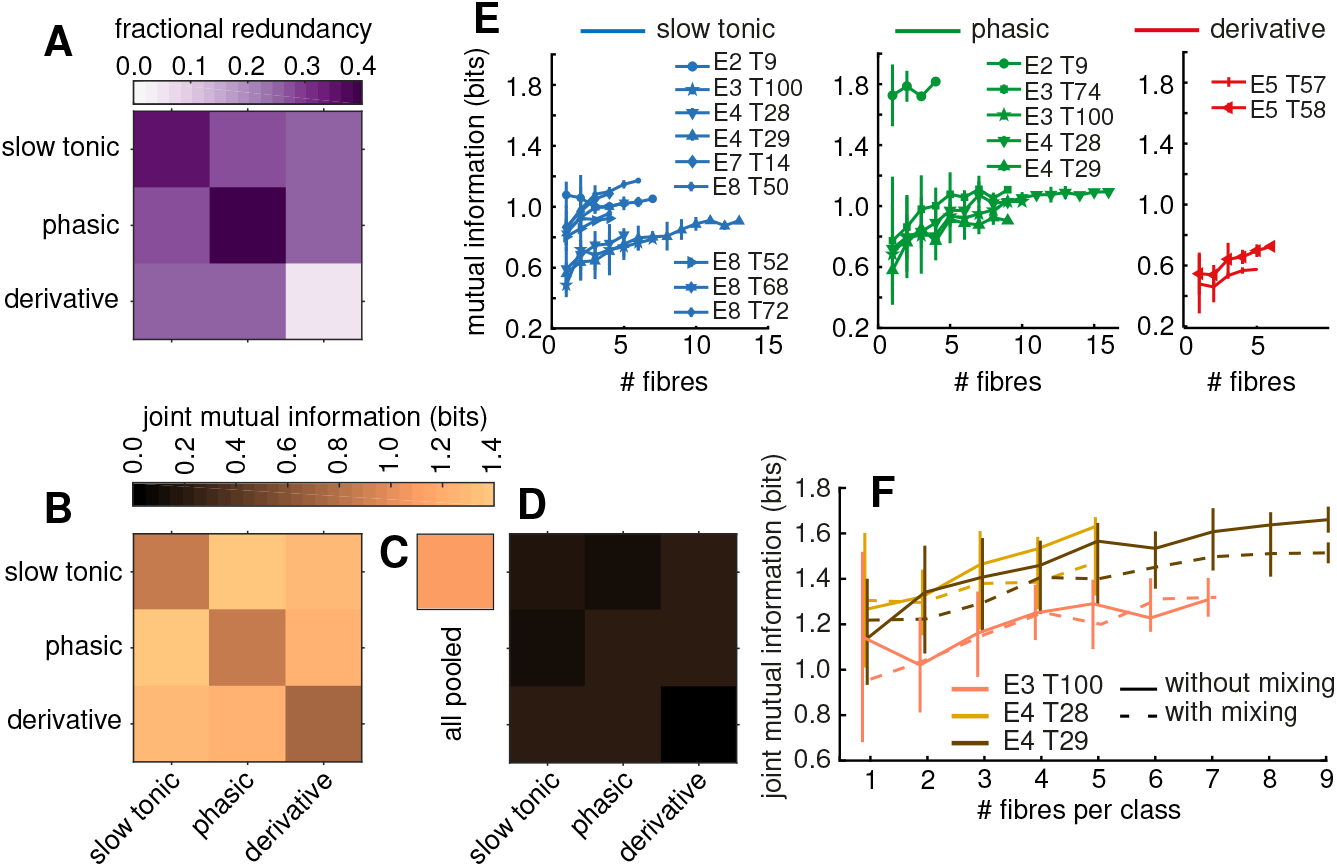
Combining complementary fibre groups leads to high joint mutual information at a moderate redundancy. Real data. **A** Fractional redundancy between average firing rates within fibre types. For on-diagonal entries, fibres were split into two equally large subpopulations of randomly chosen fibres of the same type five times between which the joint mutual information was computed and then averaged over repetitions; trials had to contain at least 4 fibres of the same type. **B** Mutual information between average firing rates within fibre type and pressure on the diagonal, averaged across trials. Joint mutual information between average firing rates of two different types and pressure in off-diagonal entries. **C** Mutual information of the average firing rate of all fibres and pressure, mean across trials. **D** Mean mutual information of single fibres across trials on the diagonal and joint mutual information of two fibres in the off-diagonal entries. **E** Mutual information between the average firing rate across an increasing number of fibres and pressure. **F** Joint mutual information between the mean firing rates of two growing pools of fibres; phasic and slow tonic. Pools either contain one fibre type (solid line) or are mixed (dashed line). Average firing rates computed from normalized firing rates.

Pooling redundant fibres increases information rate. Still, the average firing rate across all fibres of all types does not lead to the highest attainable mutual information between population activity and pressure. Even though the pooled rate of all fibres carries a higher mutual information (Fig. 5C, 1.082 ± 0.362 bits) than the average firing rate of each individual fibre type (on-diagonal in Fig. 5B, at most 0.886 ± 0.305), it is inferior to the joint mutual information of two different fibre types combined shown on the off-diagonal entries of Fig. 5B (at least 1.255 ± 0.410 bits; see Table B2 for all MI values). This effect can be understood from the low fractional redundancy ‖ between types shown in the off-diagonal entries of Fig. 5A: the firing rates of different types are almost independent of each other (fractional redundancy approximately 25%, see Table B1). It is therefore important for the transmitted information to keep the signals from different fibre types separate. To further illustrate that mutual information depends on preserving cell type-identity, Fig. 5F displays the evolution of joint mutual information between two fibre pools of increasing size while (1) only averaging within-type (solid line) and (2) mixing types to generate two inhomogeneous pools from which the average firing rate was computed. ¶ In the case of the cleanly distinguished fibre groups of Experiment 4 (see Fig. 3), the joint mutual information of the mixed populations was clearly inferior to the homogeneous populations. In Experiment 3, the difference between mixed and not-mixed was less pronounced because of the imperfect tuning of some fibres in that experiment (partly slow tonic and partly phasic at the same time). As we saw in the beginning of this section, averaging across *similar* (redundant) fibres reduces noise without signal loss. As we see now, averaging across multiple *dissimilar* (independent) fibres, on the other hand, washes out the messages of each homogeneous fibre group and destroys information. We therefore observe a coding in separate, near-independent groups.

In Fig. 6, we confirmed the benefit of different homogeneously tuned fibre pools in surrogate data (see Sec. 2.3). This additional simulation study enabled us to partially overcome the three main shortcomings of the *in vivo* data: (1) not all cell types were recorded in each trial, (2) the low- and high-frequency power of the pressure signal were correlated, and (3) the relatively short recording duration was likely to cause finite sampling biases in our estimates of information theoretic quantities. In simulation, we selected a simple idealized pressure time course consisting of a linear slope and a sinusoid of constant amplitude (period 100 s, see Fig. 6A). The different idealized responses are shown in Fig. 6A (slow) and B (fast). As an interesting observation, the intended firing rates of the phasic fibre became very similar to the derivative response for very short (1 s) decay time constants. Making use of the increased degrees of freedom of a simulation, we implemented multiple phasic fibres with different decaying constants τ (see Table 1 for its meaning). After driving an inhomogeneous Poisson spiking process at mean firing rate 20/s with the idealized rates (shown in Fig. 6A and B) and kernel-smoothing the spikes to an estimated spike rate (see Sec. 2.3 for details), the heatmap of joint mutual information in Fig. 6C could be generated. Its values were similar to the mutual information from real data in Fig. 5B but it provides a more detailed picture. Both within the fast bladder neuron types on the lower right and the slow types on the upper left, the joint mutual information stayed low at about 0.5 bits. Within the fast group, the combination of derivative and phasic fibres with intermediate decay constants (τ=30 s) reached slightly higher values as already seen in real data. Only the combination of slow and quick fibres achieved high information rates: slow tonic (and linear) and phasic fibres combined reached the highest mutual information (~ 1.1 bits). We further observed a match between the rate of decay in phasic fibres (τ=30 s) and the dominant frequency (period *T* =100 s) in the pressure signal. When increasing the sinusoid frequency, smaller values of τ reached higher mutual information and vice versa (not shown). Fractional redundancy was high within the group of slow fibres and between phasic fibres of high and medium decay constants. As fibres became less relevant to the raw pressure signal (derivative and quickly-decaying phasic fibres), fractional redundancy decreased to about zero and the expected higher values became visible when computing redundancy towards the high-frequency component of the pressure signal (see Fig. C2). Between the cleanly separated bladder neuron types of the simulated data, the off-diagonal fractional redundancies were all close to zero - fibres were truly independent. The positive effect of averaging on mutual information that we observed in real data (Fig. 5E) was confirmed in Fig. 6E where MI rises with increasing number of fibres to average over. When comparing the MI between the fibre types and different pressure variants in the subplots of Fig. 6E, linear and phasic fibres are both informative of the raw pressure, derivative and phasic provide information about the quick components, and the fit between the intended firing rate and the pressure derivative causes an exceptionally high MI for derivative fibres and a much smaller relative benefit for added derivative fibres. In Fig. 6F we repeated the analysis of Fig. 5F with a fixed number of 10 selected fibres from each population at a firing rate of 2/s each. If we kept track of the fibre identities and only average within-type, the joint mutual information of the two fibre group mean firing rates was higher than when mixing the fibres randomly into two inhomogeneous groups - averaging between fibre types destroys information.

**Figure 6:**
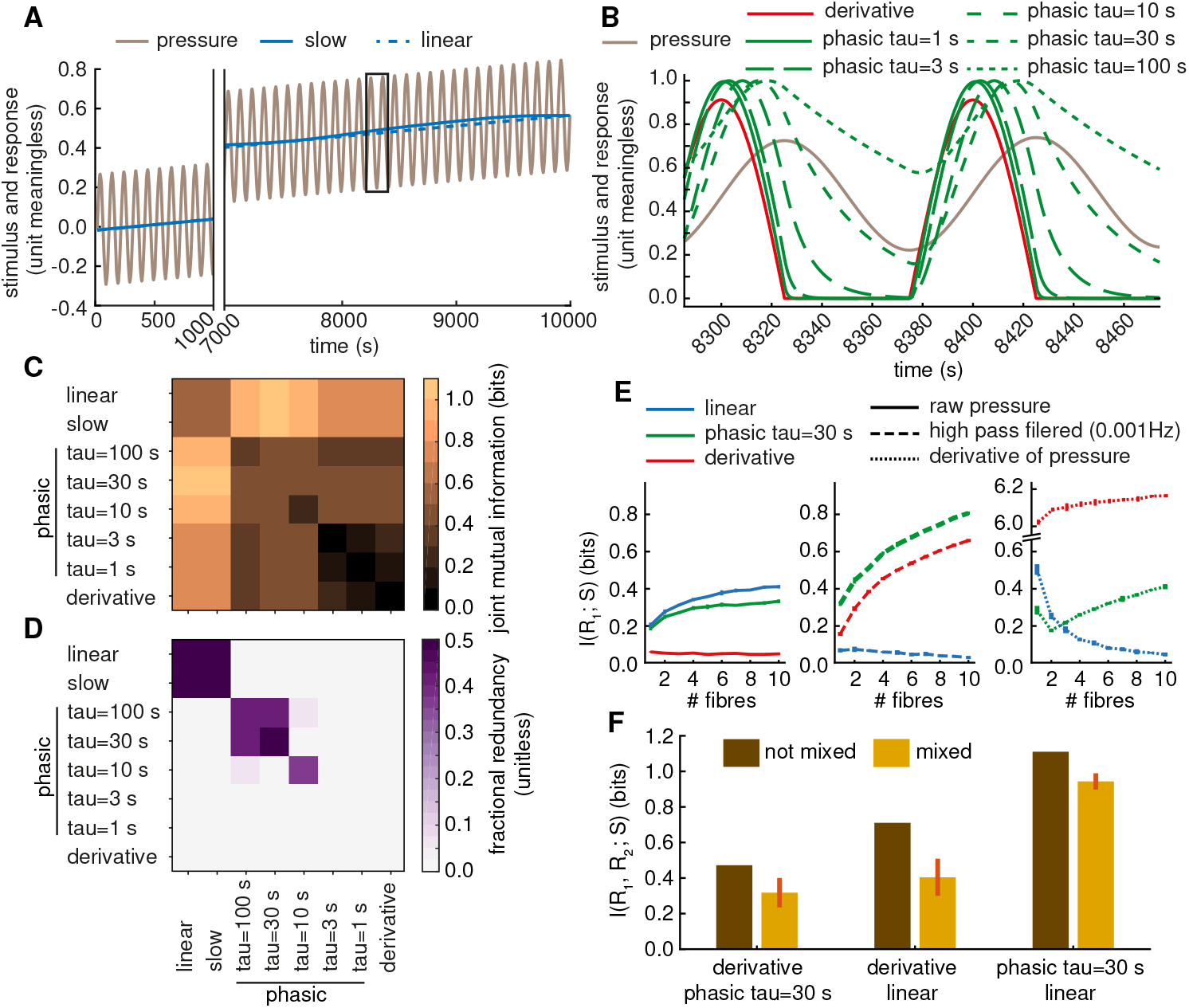
Surrogate cells confirm the benefit of complementary fibres pools on information rate. **A** Simplified pressure time course used in the simulation study along with the idealized responses of ‘slow’ and ‘linear’ fibres. **B** Idealized responses of the fast fibre types. **C** Joint mutual information between single fibres of each type in (A) and (B) with firing rate 20/s. Each square is obtained as the mean over 10 repetitions of calculating the spike times from the inhomogeneous Poisson process, kernel-smoothing for firing rate estimation and computing the single repetition joint mutual information. For on-diagonal entries, the joint mutual information between two fibres of the same type and pressure was evaluated. **D** Fractional redundancy obtained through the same process as the joint mutual information shown in (C). **E** Mutual information between an average firing rate of a population of increasing size and the raw pressure, the high-pass filtered pressure, and the pressure derivative. Firing rate of each fibre 2/s. **F** Joint mutual information of two average firing rates across 10 fibres (firing rate 2/s each) and pressure; pools either homogeneous (one fibre type) or mixed 5 times.

In summary, a partly redundant (within-type) and partly independent (between-type) coding scheme offers reliability and high SNR per channel by redundancy and a high information rate through complementary groups of bladder neurons.

### 3.3. A robust decoder based on stereotypical bladder neuron clusters

After identifying different recurring cell types by both global clustering in correlation-feature plane and by local activity clustering within trials, we demonstrated the functional significance of these groups for pressureencoding using information theory. In this last section we want to apply these encoding insights to the design of adapted decoding strategies to be used in next generation closed-loop bioelectronic medicines for bladder dysfunction. First of all, Fig. 7 shows example traces of two different trials using an OLE and a Kalman filter and different pooling methods. Single cells and pooled cell groups perform almost equally well, Kalman seems to cause less exaggerated peaks than OLE.

**Figure 7:**
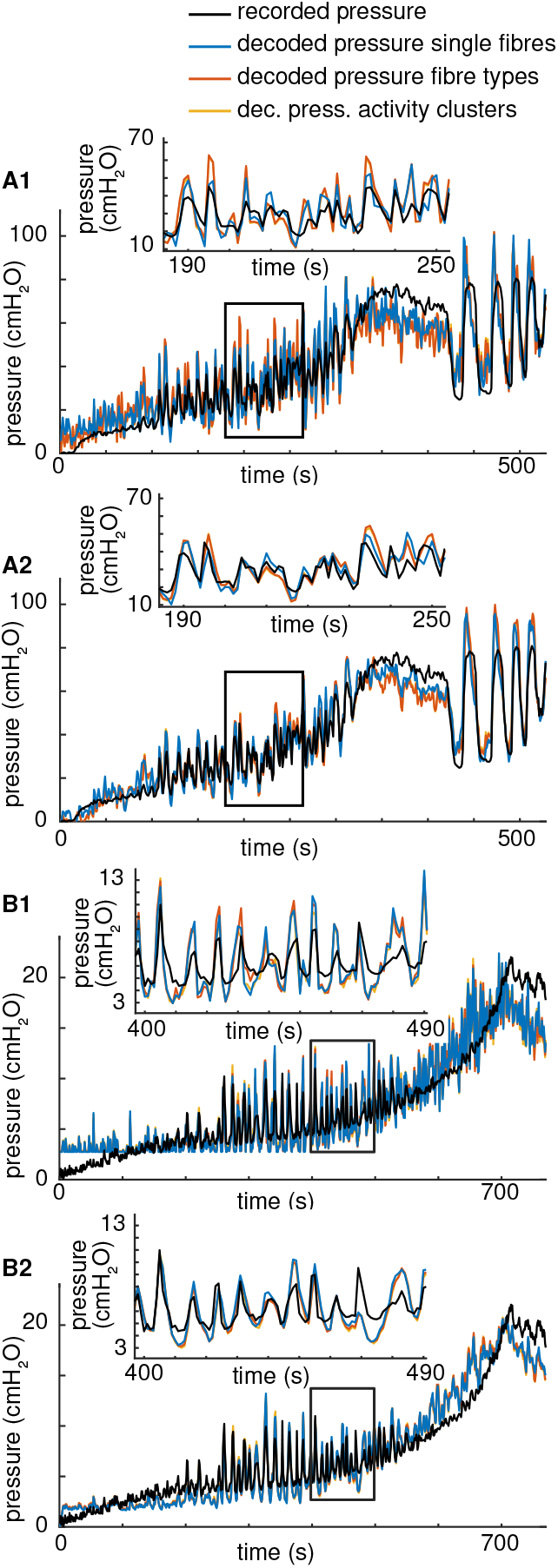
Single cells and pooled cells lead to similar decoding performances. **A1** E4 T29 decoded with an OLE. **A2** E4 T29 decoded with a Kalman filter. **B1** E8 T68 decoded with an OLE. **B2** E8 T68 decoded with a Kalman filter. Half the data points were used for training, half of them for the shown decoding.

Our information theoretic analysis showed that averaging within fibre type and keeping distinct types separate leads to a high information rate. A simple optimal linear estimator and a Kalman filter for comparison were therefore trained on both single fibres and on the mean firing rates within fibre types or activity clusters (see Table A1 for their relationship). Fig. 8 gives a visual overview of the decoding error across trials for both decoders. The bars display the 5-fold cross validated error within-trial when both training and testing on intact fibre populations. In some trials, only very few fibres were recorded and single cell firing rates were equivalent to pooled responses. This explains the low differences between single and pooled performance in experiments 1, 7, and 8. See also Table A1.

**Figure 8:**
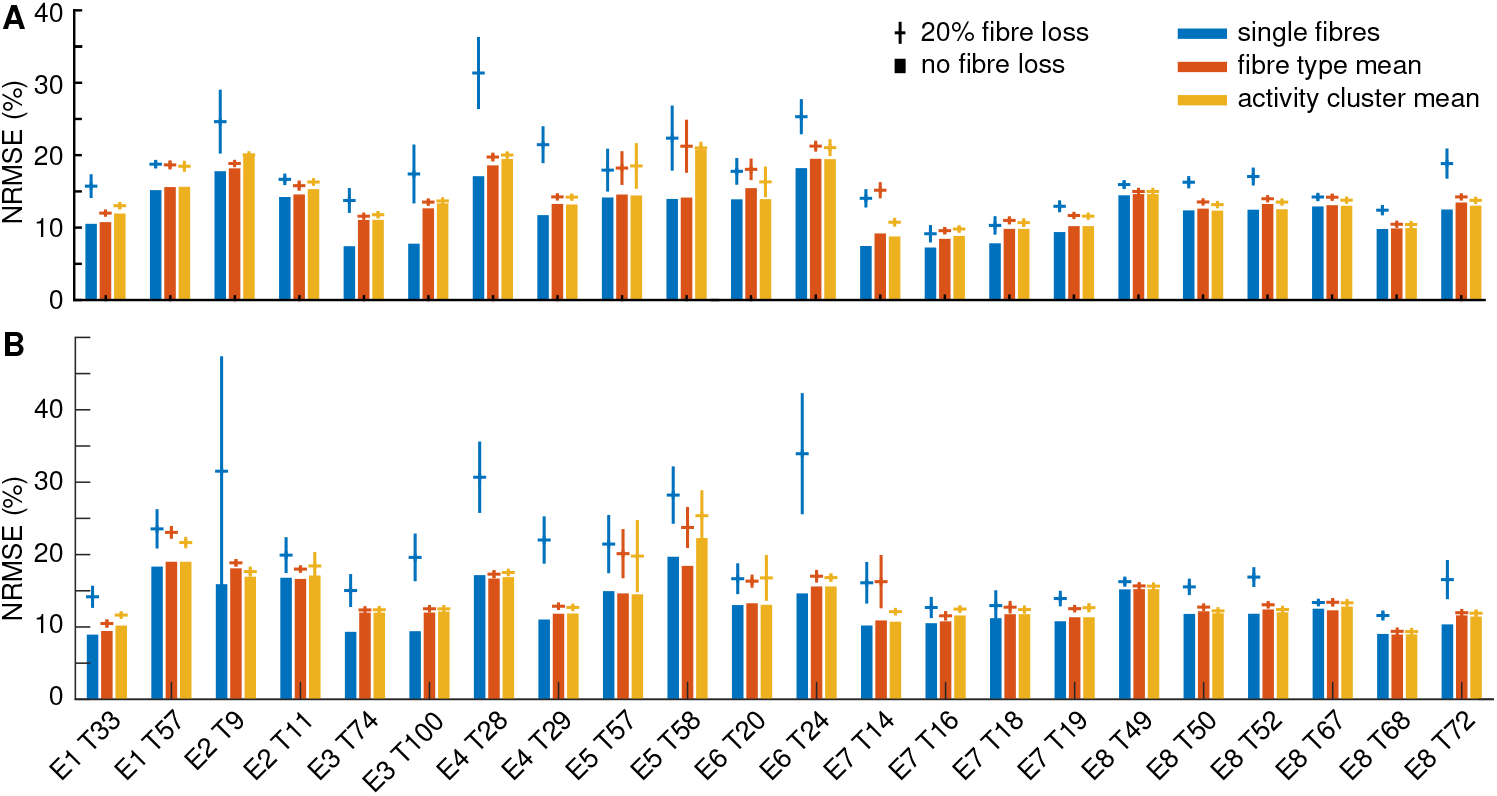
Decoding from pooled fibre types and activity clusters is robust against cell loss. **A** OLE, **B** Kalman filter. Bars show the mean decoding test error (normalized by the maximum pressure) of the decoders in a 5-fold cross validation (CV) when training and testing on the complete fibre sets. Error bars and horizontal lines indicate the standard deviation and mean test error (again mean across 5-fold CV) across 10 repetitions of removing 20% of the cells between training and testing.

For a quantitative comparison, we refer to Table 5. Values in parentheses in the following text give the values for Kalman filter, first value is for OLE. It can be seen that decoding from average firing rates (both fibre type mean and activity cluster mean) performs mildly (on average 9% (5%) and 12% (5%) higher error for fibre type and activity cluster means respectively for OLE) but significantly worse (p-values 0.00017 (0.00331) for fibre type and 0.0004 (0.00077) for activity cluster means in a paired *t*-test; 0.029 (0.058) and 0.032 (0.057) for the first trials of each experiment only), than decoding from all single cell responses. Decoding from fibre type mean firing rates tends to be marginally more successful than from activity cluster means. For comparison, we added the decoding errors for ‘electrode pools’, meaning we pooled cells per electrode (multi-unit activity, not raw threshold crossings). Performance was quite similar to the cell type and activity clusters as expected from the homogeneous electrodes, see Table 4. We test the robustness of our proposed decoding scheme by removing 20% of the cells *after training* and testing on a corrupt fibre set from which mean-responses were re-calculated. As can be seen in Fig. 8, the decoding error from single cells often became much larger after cell loss than when decoding from average responses, especially in cases like experiment 4 where many cells of each type allow for reliable cluster mean responses despite cell loss. The results are not unique to the Optimal Linear Estimator (Fig. 8A), but are very similar for a Kalman filter (Fig. 8B). The values in Table 5 confirm that the decoding error after cell loss from single cells was 18% higher than from subpopulation averages (p-values 0.00066 (0.00010) for fibre type and 0.00053 (0.00009) for activity cluster means; 0.043 (0.078) and 0.062 (0.016) for the first trials only). While for the single trials per experiment (8 observations), some comparisons are not statistically significant anymore at the 0.05 significance level, the main message prevails: redundancy leads to reliability.

**Table 5:** Decoding from pooled fibre subpopulations is more robust. Values are mean and standard deviation of NRMSE across trials.

|  | signal type | no cell loss | 20% cell loss |
| --- | --- | --- | --- |
| OLE | single fibres | $0.121 \pm 0.034$ | $0.176 \pm 0.049$ |
| | fibre type | $0.132 \pm 0.030$ | $0.149 \pm 0.036$ |
| | act. cluster | $0.136 \pm 0.035$ | $0.149 \pm 0.037$ |
| | electrode | $0.133 \pm 0.039$ | $0.155 \pm 0.045$ |
| Kalman | single fibres | $0.128 \pm 0.031$ | $0.186 \pm 0.059$ |
| | fibre type | $0.134 \pm 0.028$ | $0.150 \pm 0.038$ |
| | act. cluster | $0.135 \pm 0.030$ | $0.149 \pm 0.036$ |
| | electrode | $0.135 \pm 0.033$ | $0.160 \pm 0.058$ |

Clustering can have advantages for decoding beyond an increased robustness against cell loss. Grouping fibres periodically by their recorded activities may allow for a continuous identification of relevant cells *without knowledge of the pressure signal*. Similarly firing fibre groups are most likely driven by the same stimulus and if a subset of these similar fibres is already known to be bladder units, clustering offers an unsupervised way of identifying new relevant fibres on-line in the face of varying recording conditions caused by e.g., electrode migration.

## 4. Discussion

We have shown that bladder neurons of stereotypical response characteristics exist and assigned them to three types - slow tonic, phasic and derivative. Together, the different types implement a partly redundant, partly complementary encoding scheme for intravesical pressure that achieves a reliable and effective information transmission. We clustered fibres globally across all trials from their correlations to differently filtered variants of the pressure signal and reproduced these fibre types through unsupervised activity clustering within trials. In both real data and surrogate cell populations, we quantified the benefit of within-type redundancy (reliability, enhanced signal-to-noise ratio) and between-type tuning differences (maximization of transmitted information by complementary channels) using information theory. Building on these encoding insights, we proposed an informed decoding scheme that builds on cluster (feature-based or activity-based) mean firing rates and thereby offers increased robustness at a moderate accuracy reduction.

One limitation of our study was the sparse sampling of fibres. Using microelectrode arrays, we could record from 6 to 125 cells in each trial - of which at most 23 were identified as bladder-units. Given the high number of cell bodies in the S1 and S2 DRG of cats (~12000 [29]) and bladder-units (~1000 [30, 28, 31, 32]), we thus recorded from at most 2% of the overall bladder-unit population. This sparseness may well be the cause of the observed variability in the distribution of cell types across trials shown in Table A1 and leaves uncertain whether cell types exist in consistent ratios across animals. The study is further limited by the pressure signal that drove the bladder neurons we were analyzing. Firstly, the pressure did not contain much high frequency power, keeping us from conducting a sophisticated frequency-analysis or bladder neuron responsiveness mapping such as spike triggered averaging. Secondly, the non-stationary nature of the pressure signal and the limited reproducibility of the pressure signal across trials prevented a principled error-analysis of our information theoretic measures (e.g., by bootstrapping). Lastly, the high-frequency events (contractions) usually took place at high stationary pressure. Therefore, the firing rates of fibres responding to high-frequency events (phasic and derivative) were usually high in correlation to the slow signal components simply by correlation of slow and fast stimulus components. This made it difficult to distinguish ‘purely phasic’ and mixed phasic and slow tonic bladder neurons. The only way of overcoming these limitations is to record more data, best while enforcing isobaric and isovolumetric conditions.

Our clustering into three types of fibres is, as we state from the beginning, a deliberate simplification. As can be seen in Fig. 1B, tonic and phasic fibres do overlap in the correlation-feature space and this is at least partly due to a mixed bladder neuron tuning. It remains to be seen whether this overlap in responses is an imprecision of the bladder neuron expression that induces noise or is in fact a feature of the transmission strategy that our analysis does not acknowledge. Given the availability of more data, we could, instead of classifying cells as distinct types, assume a continuum of cell responses and infer the densities of latent parameters of a generating model. This more advanced analysis is left for future work.

The main limitation of our decoding study was the lack of controlled repetitions. We therefore emphasize that the advantages of clustering have to be confirmed on data with cells sorted across trials. Another limitation of the decoding scheme we proposed is its dependency on online spike sorting which in itself complicates the interface considerably. We did not observe clustering of the cell types within the electrode arrays beyond the degree expected by chance across experiments. Therefore, an unsorted ‘electrode-activity’ will often not provide a clean separation of fibre types (see Fig. 4 and Table 4). It is further possible that for much larger cell pools, a decoder would automatically select redundant sensors without the pre-processing step of explicitly grouping cells. It is also conceivable that the loss function of the decoder could be adapted to eg., penalize uneven fibre contributions. We hope that our approach sparks research towards better informed decoders that exploit the encoding of a quantity of interest by a population of neurons.

In addition to reliability and the benefits of averaging over imperfect sensors, other reasons for implementing multiple similar fibres are conceivable. Looking at the bladder and its feedback loop into the spinal cord as a control system, we observe that no quick control is required. The fastest events, single contractions, take seconds to tens of seconds. The peripheral nervous system can thus afford a considerable lag between intravesical pressure and the response by its higher control centers in the spinal cord an higher neural levels and can implement feedback by energetically cheap thin, slowly conducting fibres as it is observed [21, 33, 34, 35]. These thin fibres, however, do not fire at high frequencies, imposing a limit on the information rate per fibre [69]. The observed high number of thin similar fibres can therefore be viewed as the result of an energetic optimization of the information channel that ensures a sufficient information rate at an affordable lag [70].

The different groups of bladder neurons can be understood as reporting the two main components of the physiological pressure signal: the bladder (1) fills steadily at a very low rate of pressure change and (2) contracts ‘quickly’. It is not surprising that sensors for those two main signal components exist in slow fibres on the one hand and fast phasic and derivative fibres on the other hand. This mapping of bladder neuron responsiveness to signal components has been reported in many studies on nervous sensory processing, for instance as receptive fields in the visual and auditory cortex [71, 72].

Finally, many organ systems use an afferent encoding scheme based on stereotypical bladder neuron subpopulations, similar to our findings in the bladder. Phasic and tonic fibres have been reported in the colon [73, 74, 75], gall bladder [76], the lung (slowly- and rapidly-adapting sensors) [77, 78, 79, 80], similarly separate subpopulations were observed in muscle spindles [81, 82]. We hypothesize that the same benefits may have led the evolution of all these sensory populations towards an identical encoding scheme: complementary channels, each reliable due to within-type redundancy, independently encode different (quick and slow) aspects of the quantity of interest and together achieve a high information rate.

## Acknowledgement

This work was funded by the EPSRC CDT in Neurotechnology for Life and Health (EPSRC grant EP/L016737/1) and Galvani Bioelectronics; NSJ thanks EP/N014529/1 and EP/K503733/1. We thank Nikolas Barrera for spike sorting experiments 6 to 8.

## Appendix A. Detailed analysis of the relation between activity clusters and cell types

In addition to the summary of the given in Table 3, Table A1 gives a more detailed overview of the relationship between activity clusters and cell types (feature clusters) in all trials. In some trials with multiple cell types, activity clusters were inhomogeneous: E2T9, E3T100, E8T72. Other diverse trials were more successful, with each cluster capturing one specific cell type: E1T57, E2T11, E4T28, E4T29, E8T67.

---

**Table A1:**
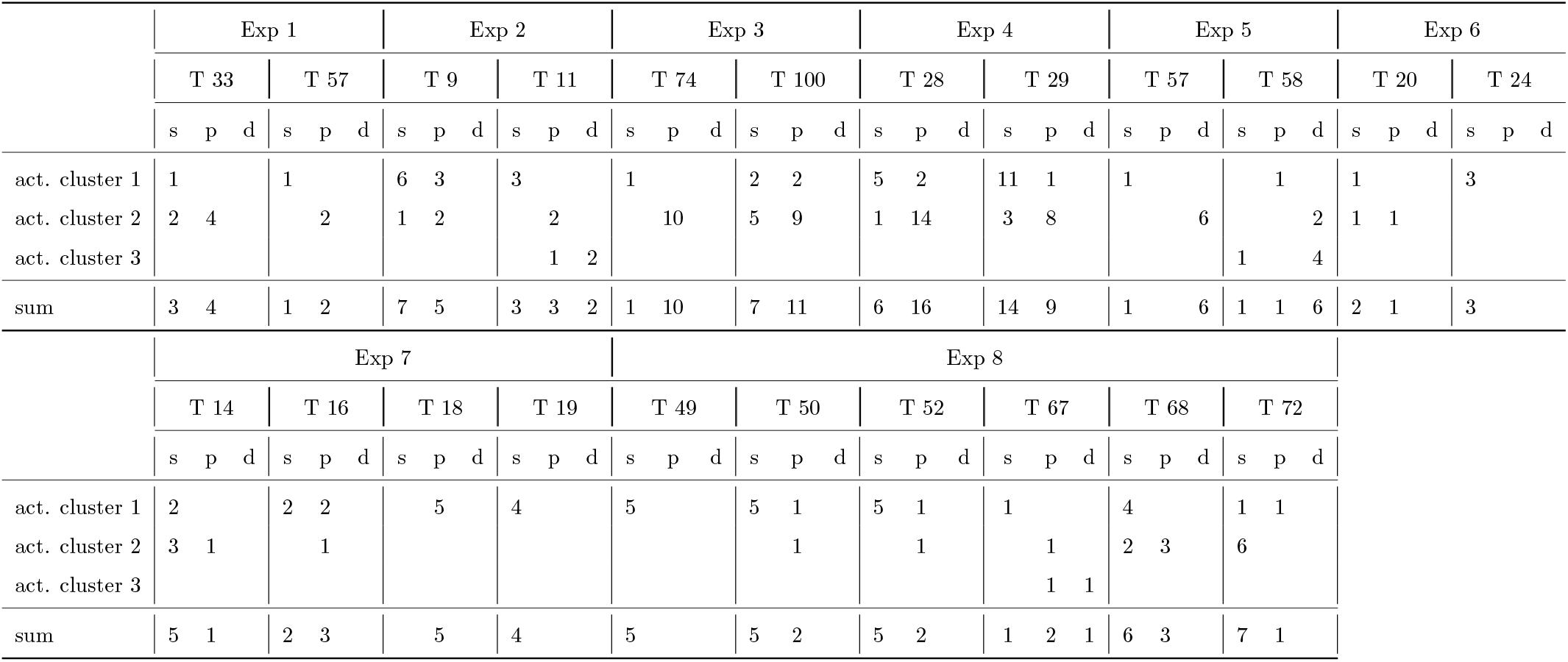
Unsupervised clustering by activity often yields near-homogeneous groups of one bladder neuron type. Each trial is shown separately and the three columns per trial correspond to the three fibre types slow tonic (s), phasic (p), and derivative (d). In each row, fibres of the three activity clusters are counted. Ideally, each activity cluster would only contain one of the three fibre types. Numbers are fibre counts.

## Appendix B. Tables of mutual information and redundancy

The following tables give the numerical values for mutual information and fractional redundancy shown in Fig. 5A, B, and D.

---

**Table B1:**
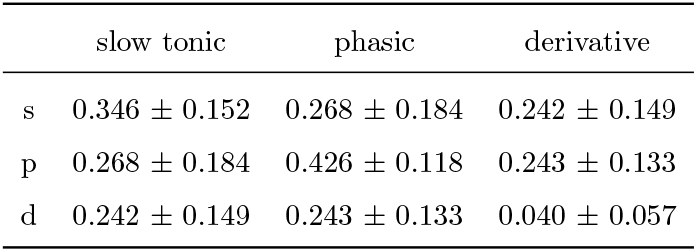
Figure 5A. Fractional redundancy between within-type mean firing rates; mean across trials. All values in bits.

**Table B2:** Figure 5B. Joint mutual information between within-type mean firing rates; mean across trials. All values in bits.

|  | slow tonic | phasic | derivative |
| --- | --- | --- | --- |
| s | $0.886 \pm 0.305$ | $1.374 \pm 0.375$ | $1.281 \pm 0.362$ |
| p | $1.374 \pm 0.375$ | $0.865 \pm 0.410$ | $1.255 \pm 0.410$ |
| d | $1.281 \pm 0.362$ | $1.255 \pm 0.410$ | $0.709 \pm 0.267$ |

**Table B3:** Figure 5D. Joint mutual information between single fibre firing rates, mean over all fibres of all trials for each type after MI calculation. All values in bits.

|  | slow tonic | phasic | derivative |
| --- | --- | --- | --- |
| s | $0.182 \pm 0.210$ | $0.137 \pm 0.177$ | $0.209 \pm 0.179$ |
| p | $0.137 \pm 0.177$ | $0.214 \pm 0.145$ | $0.199 \pm 0.147$ |
| d | $0.209 \pm 0.179$ | $0.199 \pm 0.147$ | $0.036 \pm 0.039$ |

## Appendix c. Redundancy towards fast pressure components

Fractional redundancy of two firing rates only evaluates to meaningful values when computed in relation to a relevant signal with which at least one firing rate has a high mutual information. We therefore obtain very small fractional redundancies in relation to the raw pressure signal for very quick phasic and derivative fibres. Here, we repeat the redundancy computation in relation to only the quick components of the pressure.

---

**Figure C1:**
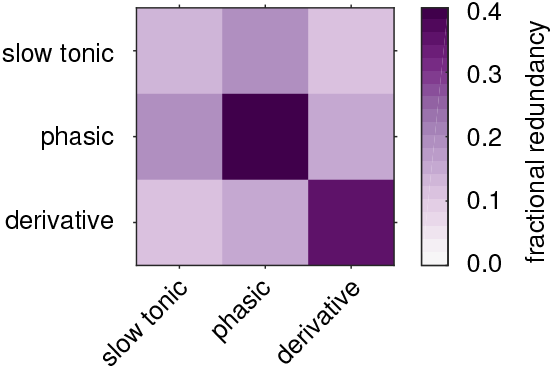
Derivative fibres only show their within-type redundancy with respect to the derivative of pressure. Real data. Fractional redundancy of fibre type average firing rates and the derivative of pressure.

**Figure C2:**
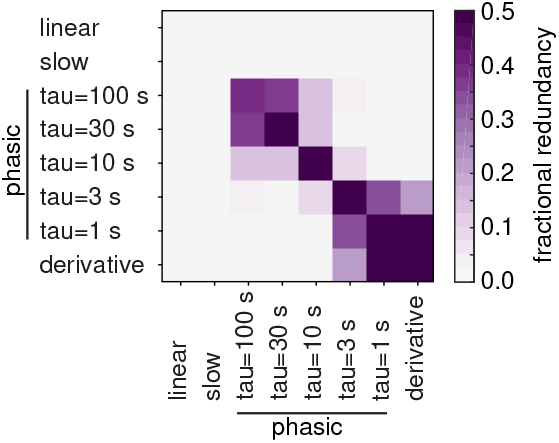
Fast fibres only show their mutual redundancy with respect to the high-pass filtered pressure. Surrogate data. Fractional redundancy of simulated firing rates and the high-pass filtered pressure above 0.001 Hz.

## Footnotes

‡ The pressure was first low-pass filtered at a high frequency of 0.25 Hz to remove noisy transients. The derivative was computed as the step-wise difference between samples of this filtered pressure.

§ As single fibre responses were often too noisy to obtain meaningful redundancy estimates, we here computed it between within-type average firing rates.

‖ Derivative fibres are by themselves not very informative of the raw pressure signal and their within-type fractional redundancy becomes less meaningful. If we compute redundancy relative to the derivative of the pressure signal as shown in Fig. C1, fractional redundancy also reaches high values for this fibre type.

¶ The shown example trials were chosen because they had at least 4 slow tonic and 4 phasic fibres.

